# Computational image analysis reveals the structural complexity of *Toxoplasma gondii* tissue cysts

**DOI:** 10.1101/2020.05.21.108118

**Authors:** Neda Bauman, Andjelija Ilić, Olivera Lijeskić, Aleksandra Uzelac, Ivana Klun, Jelena Srbljanović, Vladimir Ćirković, Branko Bobić, Tijana Štajner, Olgica Djurković-Djaković

## Abstract

*Toxoplasma gondii* is an obligate intracellular parasite infecting up to one third of the human population. The central event in the pathogenesis of toxoplasmosis is the conversion of tachyzoites into encysted bradyzoites. A novel approach to analyze the structure of *in vivo*-derived tissue cysts may be the increasingly used computational image analysis. The objective of this study was to quantify the geometrical complexity of *T. gondii* cysts by morphological, particle, and fractal analysis, as well as to determine if and how it is impacted by parasite strain, cyst age, and host factors. Analyses were performed on 31 images of *T. gondii* brain cysts of four type-2 strains (the reference Me49 strain and three local isolates, named BGD1, BGD14, and BGD26) using ImageJ software package. The parameters of interest included diameter, circularity, relative particle count (*RPC*), fractal dimension (*FD*), lacunarity, and packing density (*PD*). Although cyst diameter varied widely, its negative correlation with *RPC* was observed. Circularity was remarkably close to 1, indicating that the shape of the brain cysts was a perfect circle. *RPC*, *FD,* and *PD* did not vary among cysts of different strains, age, and derived from mice of different genetic background. Conversely, lacunarity, which is a measure of heterogeneity, was significantly lower for BGD1 strain vs. all other strains, and higher for Me49 vs. BGD14 and BGD26, but did not differ among Me49 cysts of different age, and derived from genetically different mice. This study is the first application of fractal analysis in describing the structural complexity of *T. gondii* cysts. Despite all the differences among the analyzed cysts, most parameters remained conserved. Fractal analysis is a novel and widely accessible approach, which along with particle analysis may be applied to gain further insight into *T. gondii* cyst morphology.

## Introduction

*Toxoplasma gondii* is an obligate intracellular parasite belonging to the phylum Apicomplexa that can infect all warm-blooded animals, including humans. It is considered as one of the most successful parasites on the Earth, estimated to infect up to one third of the human population [1]. While the infection is generally mild and self-limiting in immunocompetent individuals, it has the potential to cause severe outcomes in immunocompromised patients and the fetus [2].

The three infectious life stages of *T. gondii* are rapidly proliferating tachyzoites, slowly growing bradyzoites (inside tissue cysts), and sporozoites (inside oocysts). During its complex life cycle, tachyzoites differentiate into bradyzoites, which in an immunocompetent host form tissue cysts, predominantly in the central nervous system and muscles [3]. Encystment is recognized as a central event in the pathogenesis, persistence, and transmission of toxoplasmosis.

Studies focusing on tissue cyst biology have been conducted for decades. Despite numerous *in vitro* studies [4, 5, 6], *in vivo* research that may provide more reliable observations on the biology of *T. gondii* cysts is still limited [7].

In recent years, different types of computational image analysis have found their application in life sciences, with the potential to change the future of medicine [8, 9]. Fractal analysis, as a method for quantifying the complexity of natural objects, is being increasingly used in biomedical research [10, 11, 12, 13], including a few studies in the field of microbiology, focusing on bacteria [14, 15], fungi [16], and *Plasmodium* [17]. In addition, particle analysis, a computational biology method for precise cell counting [18], has been successfully applied in several studies for the counting of parasites, including *T. gondii* [7, 19]. However, fractal analysis has not yet been used for studying *T. gondii*. The ability of fractal analysis to describe natural processes in various tissues implies that it may be used for studying the structural complexity of *T.gondii* and eventually facilitate research on *in vivo*-derived tissue cysts.

This study aimed to quantify the geometrical complexity of *T.gondii* cysts by fractal analysis and to determine if and how it is impacted by a number of parasite or host factors. Additionally, morphological and particle analyses of *T. gondii* cysts were applied to gain further insight into the cyst shape uniformity, as well as to a possible correlation between the cyst size and the number of parasites.

## Material and methods

### Study design

Images of *T. gondii* cysts analyzed in this paper were obtained in two ways, by selection from the photo archive of strains isolated and maintained in the Serbian National Reference Laboratory for Toxoplasmosis (NRLT), and from cysts harvested from mice infected specifically for this study.

Four different *T. gondii* strains belonging to genotype-2 were included.

### Archived cyst image source

Three out of the four analyzed strains, referred to as BGD1, BGD14, and BGD26, were isolated from human biological materials and thereafter maintained by regular passages in Swiss Webster (SW) mice by intraoesophageal gavage of brain suspensions containing cysts of the respective strains [20, 21, 22]. In this study, archived images of cysts obtained from the 8^th^ (BGD1), 21^st^ (BGD14), and 2^nd^ (BGD26) passage, were used. These cysts were more mature than those from experiment, obtained from mice infected 6 to 15 months previously.

### Cysts obtained from experimental infection

For experimental infections, two genetically different hosts, SW and BALB/c mice (Medical Military Academy Animal Research Facility, Belgrade, Serbia) were used. All mice were female and weighed 18-20 g at the beginning of the experiment. Mice were housed five per cage at the Institute for Medical Research Animal Research Facility, and offered regular mouse feed and drinking water *ad libitum*.

Mice were inoculated intraperitoneally (i.p.) with 10^4^ tachyzoites of the reference Me49 strain and the BGD1 strain (from the 32^nd^ passage). Tachyzoites were harvested from cultivated Vero (ATCC No. CCL-81) cells inoculated with peritoneal exudates of mice, previously infected i.p. with cysts. Me49 parasites were used for infecting both SW and BALB/c mice and cysts were harvested at two time points, at 6 and 12 weeks post-infection (p.i.). BGD1 parasites were inoculated only into SW mice, and cysts were harvested 8 weeks p.i.

To harvest brain cysts, mice were sacrificed by cervical dislocation, brains removed and homogenized with a syringe in 1mL of saline each.

### Visualization of *T. gondii* cysts

For cyst visualization, 25 μl of the brain suspension was placed on slides and examined under a phase-contrast microscope (Axioskop 2 Plus Zeiss) magnification of 1000x; cysts were morphologically recognized and photographed by Zeiss AxioCamMRc 5. Diameter was measured immediately after taking the picture of microscopically visualized cyst. Representative images were selected for further analysis.

### Circularity and particle analysis

Images of *T. gondii* cysts, saved as the grayscale TIFF, were studied by means of morphological, particle, and fractal analysis. All analyses were performed using the ImageJ software package (NIH, Bethesda, MD, USA). The main morphological parameter [23, 24] of interest was cyst circularity, defined as

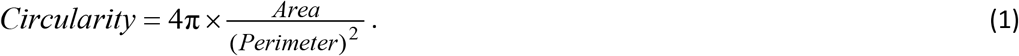

Circularity is a measure of how close a shape is to a perfect circle. Shapes close to a circle have circularity values close to 1.0, whereas elongated shapes have very low circularity (close to 0.0). Cysts were carefully manually delineated, as shown in Fig 1A. Parameters including *Area*, *Perimeter*, and *Circularity* were accurately determined by the ImageJ software. The delineated cyst cross-sectional areas were transferred to black background images for subsequent particle analysis.

**Fig 1.**
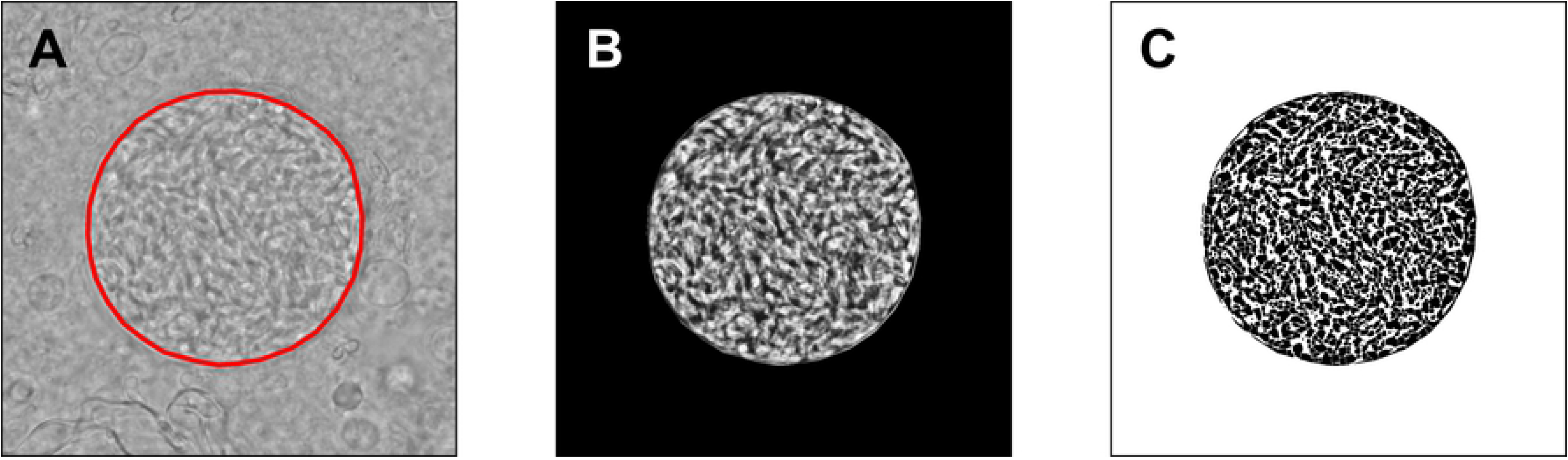
Circularity and particle analysis of *T. gondii* cysts. (A) - A cyst contour was precisely delineated and the closeness of this contour to a perfect circle was quantified as Circularity, a parameter accurately calculated by the ImageJ software. (B) - A contrast-enhanced cyst cross-sectional area, following the histogram equalization, was used in subsequent thresholding and calculation of parameters. (C) - The relative particle count (*RPC*) was determined following the local Bernsen auto-thresholding, watershed segmentation, and particle counting. An Me49 6-weeks-old cyst shown for illustration.

For particle analysis, images were first enhanced using histogram equalization. During this operation, the distribution of the most frequent pixel intensity values was spread out, resulting in the global contrast enhancement. The resulting image, corresponding to the original one in Fig 1A, is shown in Fig 1B. These enhanced contrast images were subjected to a local auto-thresholding, using the Bernsen method with a window radius equal to 7 pixels. The Bernsen method belongs to a group of locally adaptive thresholding methods based on contrast. The ImageJ implementation uses circular windows, which is convenient for the analysis of *T. gondii* cysts. The obtained binary (black and white, B/W) images were further segmented by a watershed algorithm, as shown in Fig 1C, to divide the larger foreground areas of black pixels into the particles to be counted. Segmentation by a watershed algorithm is typically used to enable precise counting of closely packed cells [18, 19].

It should be noted that the light areas corresponding to bradyzoites, visible in the cyst cross-section in Fig 1B, were shown as foreground black pixels in Fig 1C. These foreground pixel areas were further segmented by a watershed algorithm for two reasons: a) it seems that, due to a low local border contrast, multiple bradyzoite areas were recognized as a single entity by Bernsen thresholding, and b) the particle counting procedure, currently available in ImageJ, is known to work properly only with clearly separated particle areas. In that manner, all of the cyst cross-section images were subjected to an automated processing and the obtained particle numbers, *N*_p_, were expected to be proportional to the real numbers of bradyzoites, with very close proportionality coefficients. The parameter of interest, expected to be proportional to the actual count, was termed the ‘relative particle count’ (*RPC*), and calculated as the number of counted particles divided by the cross-sectional area. The corresponding unit is μm^−2^.

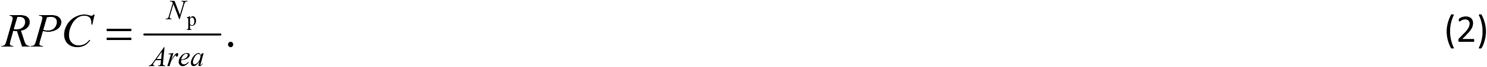

As it is very difficult to claim with certainty where the actual bradyzoite contours are located in an image, the application of the above methods allowed for a highly objective comparison of analyzed specimens. Namely, carrying out the identical procedures in all of the considered cases is expected to result in very similar proportionality coefficients with respect to an actual number of particles.

### Fractal analysis

In order to perform fractal analysis in a consistent way for all examined cysts, we prepared cutouts sized 450 pixels by 450 pixels from the central parts of all cyst cross-sectional areas, as shown in Fig 2A. The corresponding areas of the specimens equaled 580.325 μm^2^. Binary images for fractal analysis were obtained by global thresholding using two close threshold values in the vicinity of the Otsu threshold, which is equal to the average of mean grayscale levels of the two sets of pixels being partitioned. Fractal dimension (*FD*) and lacunarity of the extracted contours of the bradyzoite areas, shown in Fig 2B, were calculated by the box-counting method and cumulative mass method. Both methods calculated the *FD* with high accuracy and gave the same results.

**Fig 2.**
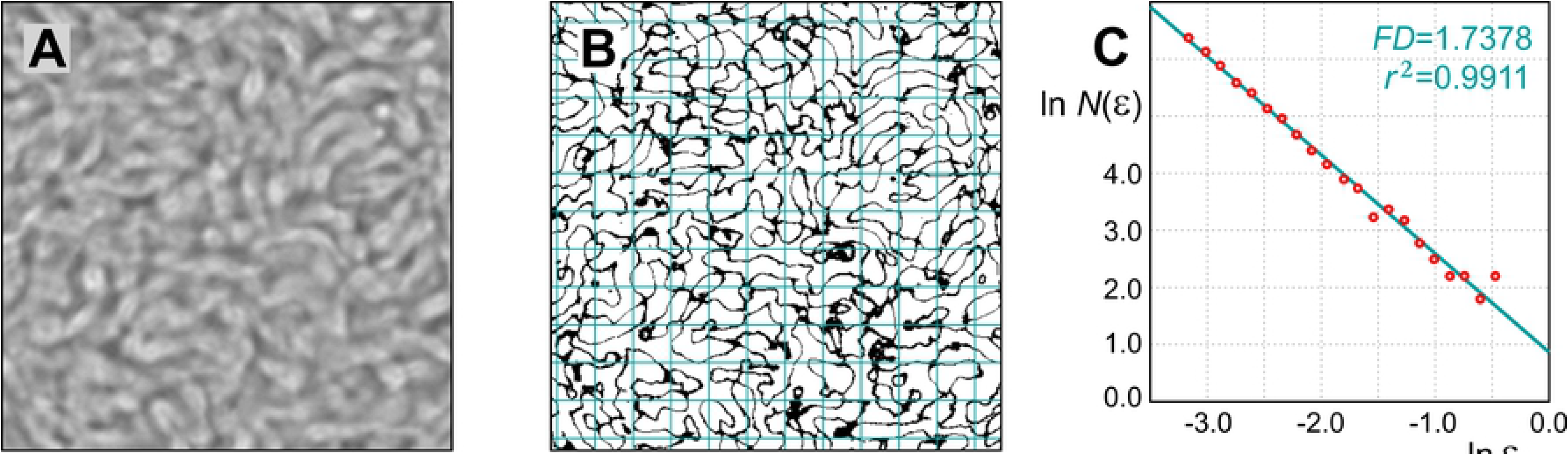
Fractal analysis of *T. gondii* cysts. (A) - Cutouts sized 450-by-450 px^2^, belonging to the central parts of cyst cross-sectional areas, were used. (B) - Corresponding binary contour images were covered by a number of grid cells of varying sizes, in order to determine fractal parameters by the box-counting and cumulative mass methods. (C) - Fractal dimensions (*FD*) correspond to the slopes of the best-fit regression lines. Each *FD* calculation is accompanied by the correlation coefficient, *r*^2^, describing a goodness of the regression line fit.

The relationship between a considered measure *η*(*l*) and a measurement scale, *l*, corresponding to the power scaling laws typical for multiplicative processes is known to be of a following form [8, 25, 26]:

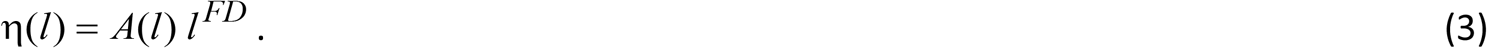

Both calculation methods rely on producing the logarithm-logarithm plot of a measure versus the scale or grid cell size and fitting the regression line based on the obtained points. We generated a series of grid cell sizes using the scaled series 7/8 (seven over eight), meaning that the ratio of consecutive grid cell sizes is about 0.875. The following grid cell sizes were used: {17, 19, 22, 25, 29, 33, 38, 43, 49, 56, 64, 74, 84, 96, 110, 126, 144, 164, 188, 214, 245, 280, 320} pixels. To reduce the calculation variability due to a particular grid cell positioning, twelve randomly positioned grids per cell size per image were used. An example positioning of a grid of cell size 38 pixels is shown in Fig 2B. In the box-counting method, *FD* is estimated from a change in the number of non-empty grid cells, *N*(*ε*) ~ *ε*^*FD*^, with a change of the cell size normalized with respect to the image perimeter *L*, *ε* = *l* / *L*, as illustrated in Fig 2C. The *FD* was obtained as a negative slope of the best-fit regression line:

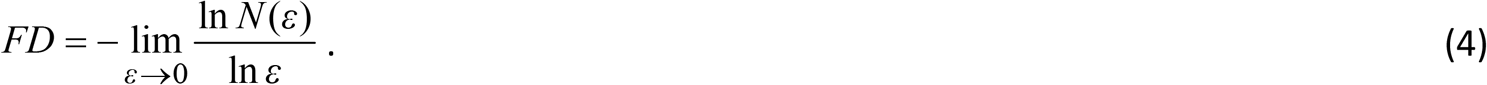

Each *FD* calculation is accompanied by the correlation coefficient, *r*^2^, describing a goodness of fit of the regression line, in a particular case.

In the cumulative mass method, a probability of finding a certain pixel number within a given cell was estimated by counting the number of pixels contained in each grid cell [25]. In this case, a positive slope of the best-fit regression line was used. The probabilities, *P*(*m*, *ε*), of finding *m* points within a cell of scale *ε*, were normalized and used to obtain the first order and higher-order mass moments:

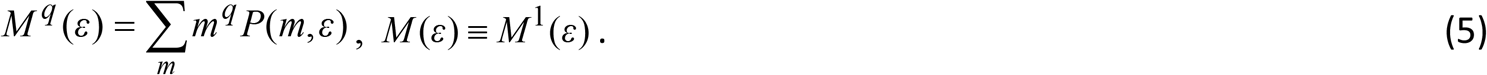

Lacunarity of a fractal set represents the ratio of the second-order moment (variance) to the first-order moment (average), of the mass probability distribution. It is a measure of structural variance within an object. Higher lacunarity is related to a larger vertical spread of the regression lines corresponding to the multiple *FD* calculations. Denoting the mathematical expectation (average)by〈·〉, the fractal lacunarity was calculated as [25, 26]:

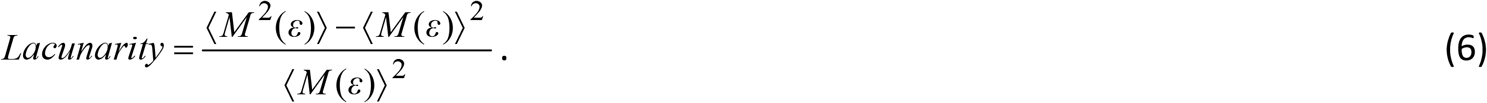

In addition to the mentioned parameters, we defined another parameter measuring the percentage of space covered by bradyzoites, and termed it ‘packing density’ (*PD*). The two-dimensional analysis was performed on the cyst cross-sectional areas captured on microscope. Both particle analysis and fractal analysis provided the number of black pixels in a considered cross section, *N*_B_. The total number of pixels in a cross section, *N*_T_, was known in all cases, which allowed the packing density estimation as

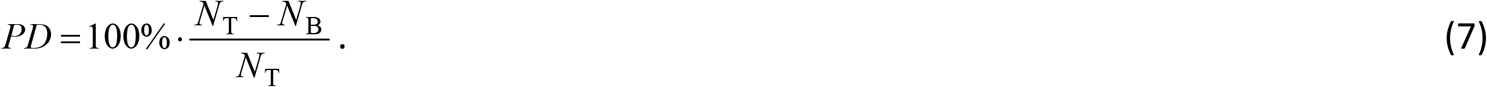

The *RPC* and *PD* parameters quantitatively estimate the number of bradyzoites in a cross section.

### Statistical analysis

Statistical analysis was performed using IBM SPSS Statistics for Windows, version 26 (IBM Corp., Armonk, N.Y., USA). The normality of the dataset was tested using Shapiro-Wilk and Kolmogorov-Smirnov tests, as well as Q-Q plots. Subsequently, image data were evaluated by analysis of variance (ANOVA) and Tukey’s test for post-hoc analysis. In cases where the Levene’s test showed unequal variances, we used Welch’s ANOVA. Independent samples t-test was used when comparing two sets of data, as well as Mann-Whitney U for comparing data with non-normal distribution. For analysis of correlation Spearman’s rank correlation test was used. The level of significance was p ≤ 0.05.

### Ethics statement

The study protocol was approved by the State Ethics Committee (Veterinary Directorate of the Ministry of Agriculture, Forestry and Water Management of Serbia decision no. 323-07-02446/2014-05/1 and 323-07-05638/2019-05). The animal experiments were conducted concordant to procedures described in the National Institutes of Health Guide for Care and Use of Laboratory Animals (Washington, DC, USA). All efforts were made to minimize suffering.

## Results

We analyzed images of *T. gondii* cysts obtained from brain homogenates of mice infected with four different parasite strains (BGD1, BGD14, BGD26, Me49) in the same host (SW); Me49 cysts obtained at two different time points (6 and 12 weeks) in the same host (BALB/c); and Me49 cysts obtained from two different hosts (BALB/c and SW). Accordingly, the analyzed cysts differed according to the parasite strain, duration of infection (cyst age), route of infection, life stage of the parasite used for infection, and host. The total number of evaluated cysts was 31. The basic properties of the source cysts are presented in Table 1.

**Table 1.**
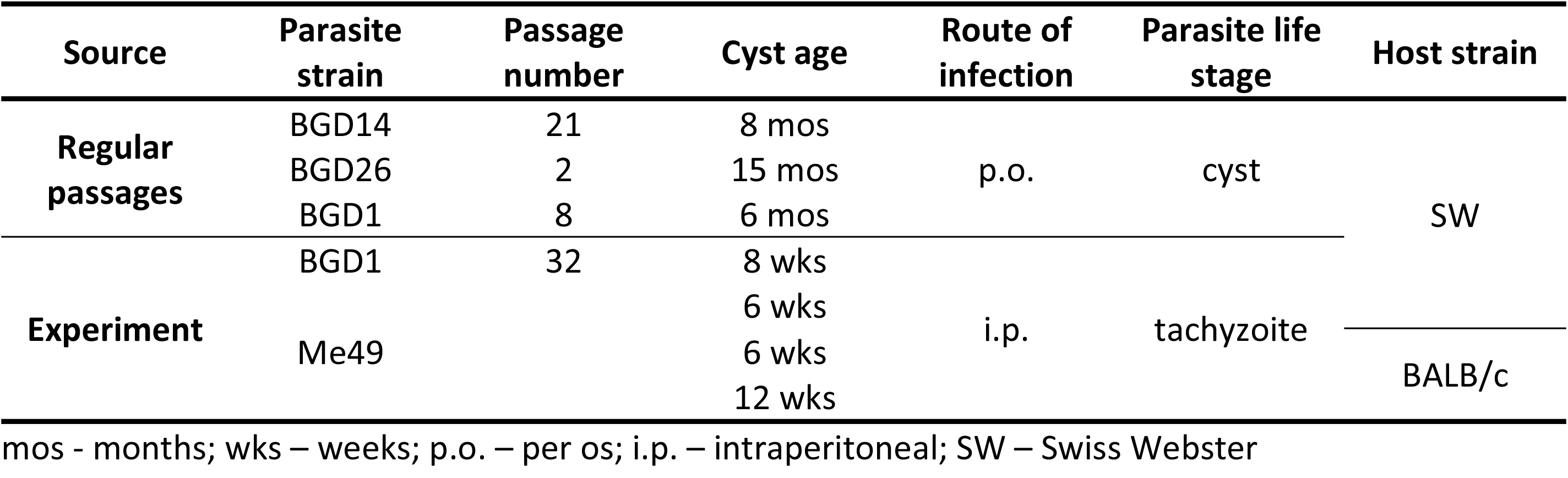
Basic properties of *T. gondii* source cysts.

Parameters of interest included diameter, circularity, *RPC*, *FD*, lacunarity, and *PD*. A summary of all results is presented in Table 2.

**Table 2.**
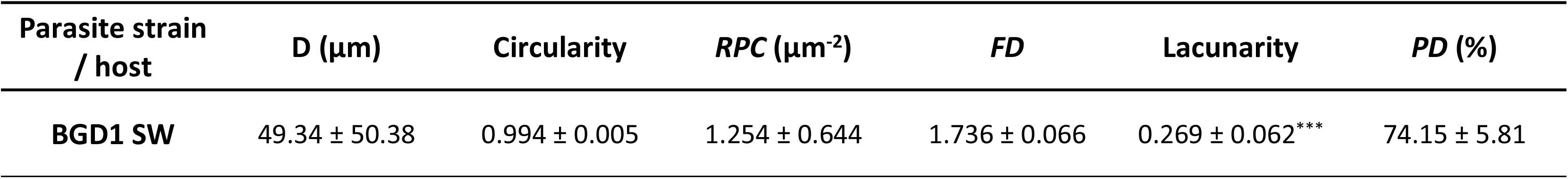
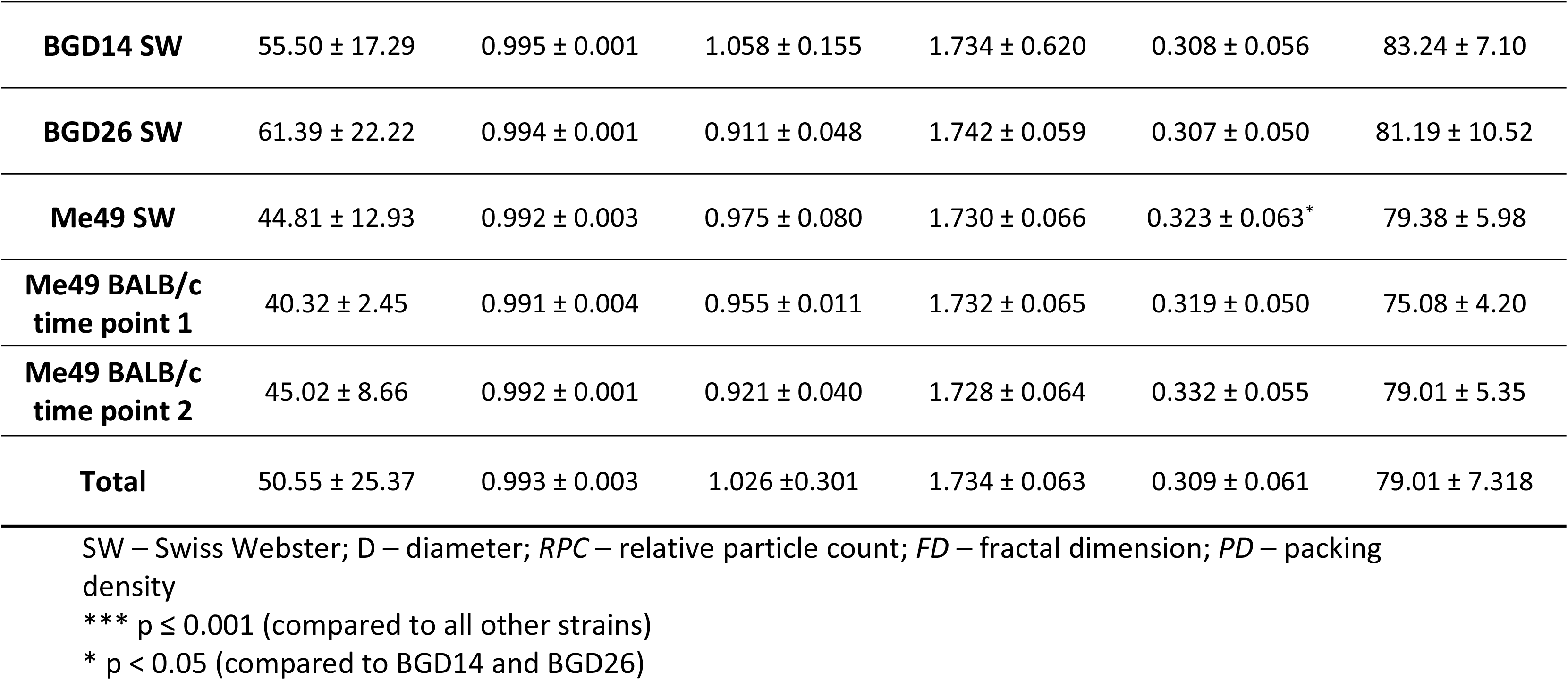
Summary of parameters of structural complexity in *T. gondii* type-2 cysts (n=31). All values are expressed as mean ± SD.

The diameter of the analyzed cysts varied widely, spanning from 12.91 μm to 132.64 μm (mean 50.55 ± SD 25.37). However, circularity was very uniform, being remarkably close to 1 for all cysts (0.993 ±0.003). Also, *PD* and *RPC* showed no statistically significant differences among cysts of different strains (p_*PD*_=0.18; p_*RPC*_=0.12), time points (p_*PD*_=0.37; p_*RPC*_=0.23), and hosts (p_*PD*_=0.46; p_*RPC*_=0.31). However, a significant negative correlation between *RPC* and cyst diameter was observed (ρ=−0.54, p=0.002).

Fractal analysis yielded interesting results. The *FD* was also very uniform among all cysts, ranging from 1.59-1.82 (mean 1.734 ± 0.063) with a variation coefficient of 0.0365 (3.6%). No difference in the *FD* was found between cysts of different strains (p=0.72), time points (p=0.51), and hosts (p=0.57) (Fig 3A; Fig 4A,B). Lacunarity, on the other hand, differed among the strains, in that it was significantly lower in the BGD1 strain in comparison to the three other ones (p≤0.001), as well as in BGD14 and BGD26 vs. Me49 (Tukey p=0.044 and p=0.033, respectively), but not between BGD14 and BGD26 (p=0.99) (Fig 3B). Moreover, it did not differ among Me49 cysts harvested at different time points (p=0.23), nor among those derived from different mouse strains (p=0.56) (Fig 4C,D).

**Fig 3.**
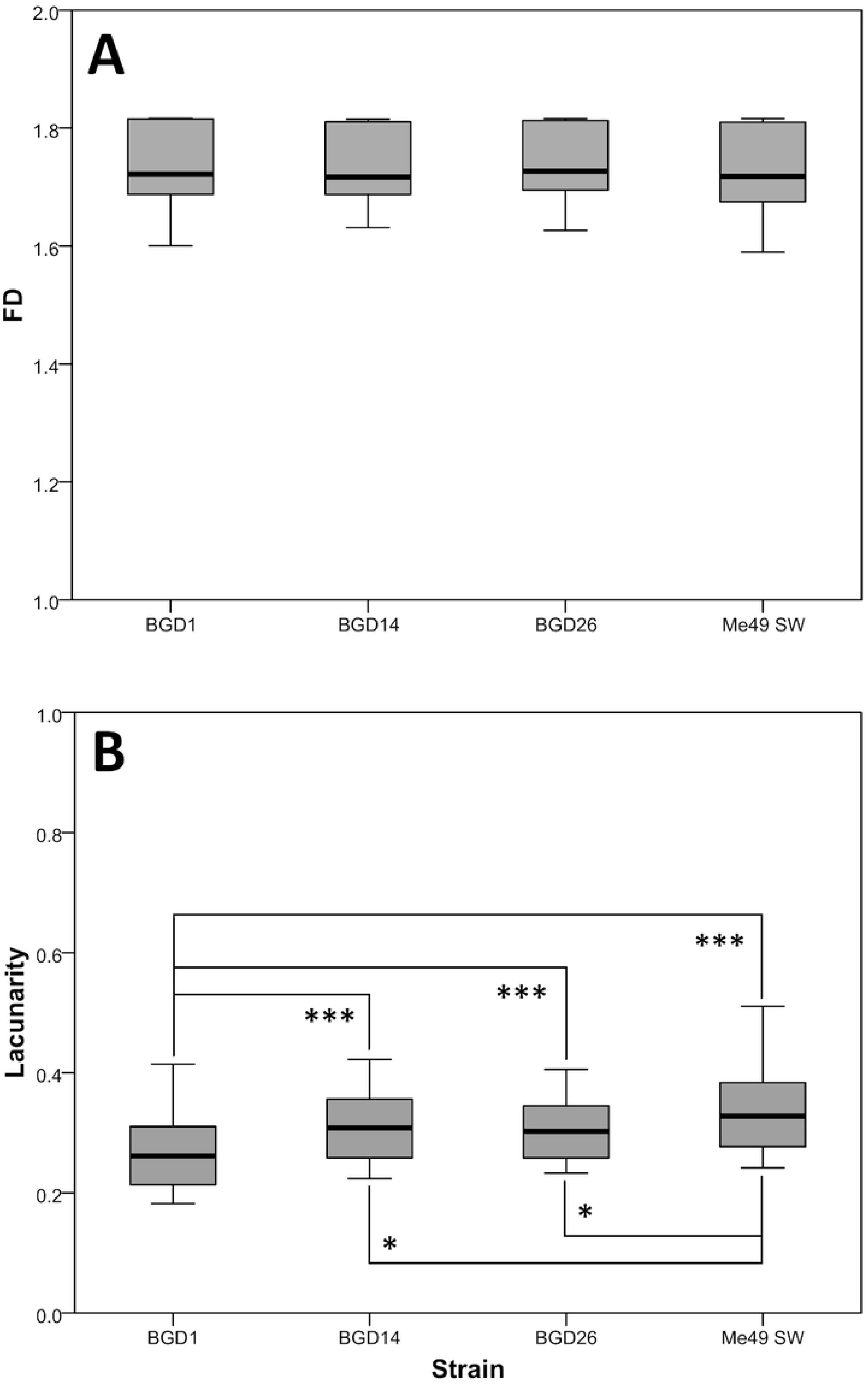
Fractal dimension (*FD*) (A) and lacunarity (B) of cysts of four *T. gondii* type-2 strains. Values are expressed as mean ± SD. (A) No statistically significant differences among the parasite strains (*F*_3,296_=0.44, p>0.05). (B) Significantly lower for BGD1 vs. all other strains, and higher for Me49 vs. BGD14 and BGD26 (*F*_3,296_=15.7, p<0.001) ***p ≤ 0.001; * p < 0.05.

**Fig 4.**
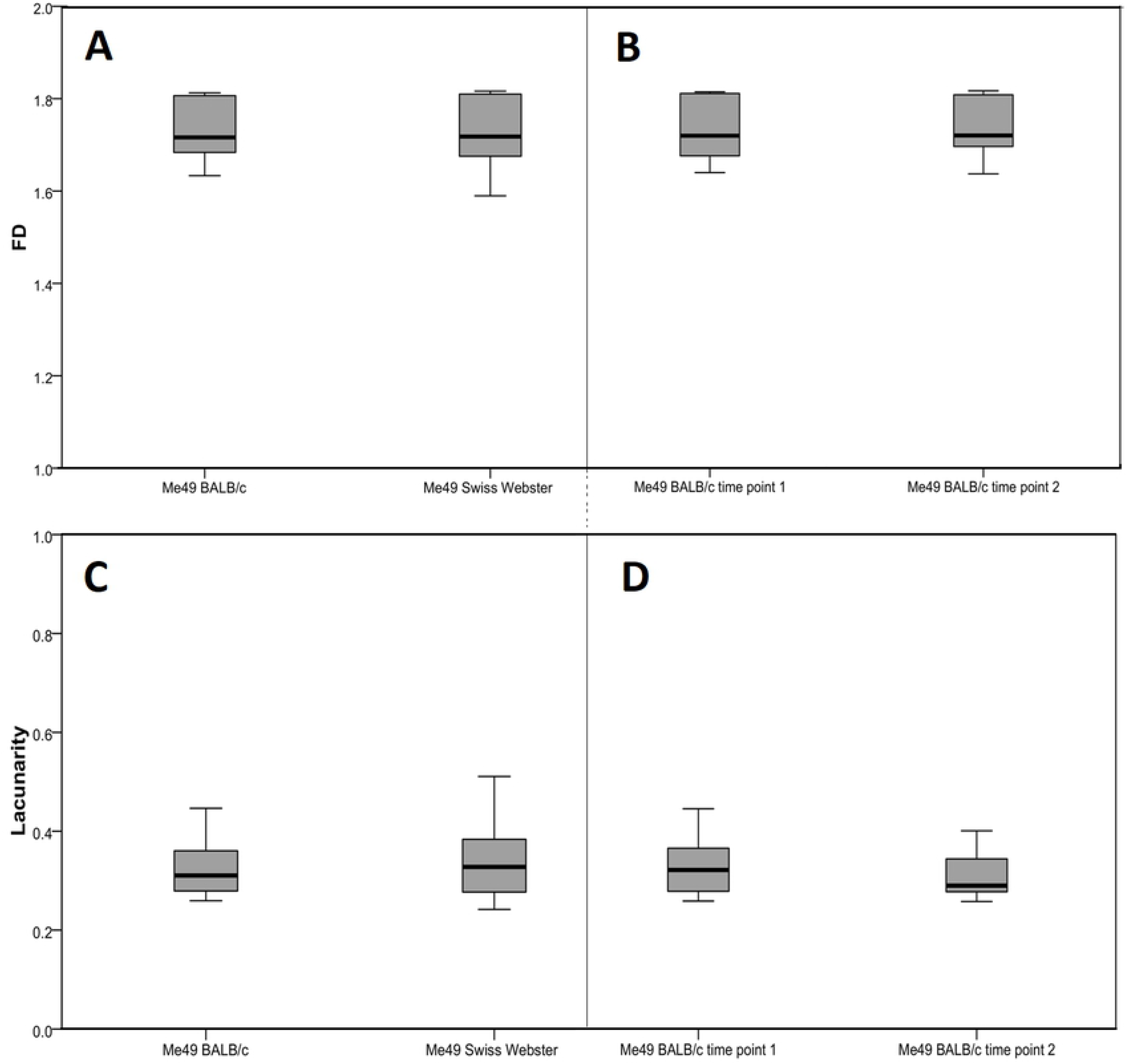
Fractal dimensions (*FD*) and lacunarity of cysts of the Me49 *T. gondii* strain according to host and cyst age. Values are expressed as mean ± SD. (A) *FD* and (C) lacunarity in BALB/c and Swiss Webster mice. (B) *FD* and (D) lacunarity at two time points (6 and 12 weeks) in the BALB/c host. No statistically significant differences were found between any two examined groups (host: *U_FD_*=2823.5, *U_lacunarity_*=2818; cyst age: *U_FD_*=573, *U_lacunarity_*=525; p > 0.05).

## Discussion

Despite years of research, questions regarding the biology of *T. gondii* tissue cysts, such as bradyzoite organization and burden within cysts, remain unresolved. An available approach to describe and quantify a structural complexity of cells and tissues is by performing fractal analysis. The growth, branching, and rearrangement of biological entities can be considered and described as a multiplicative process. The resulting structural patterns seen under a microscope possess a certain degree of space-filling and exhibit self-similarity in comparison with other areas of the same specimen, as well as with other specimens of the same type.

In an attempt to improve the understanding of the nature of the tissue cyst structural complexity, we applied fractal analysis, as well as morphological and particle analysis, to *T. gondii* cysts obtained from different sources. We set out to analyze whether tissue cyst fractal geometry values are stable or tend to change depending on the parasite strain, cyst age, or mouse host genetic background. The common feature for all of the cysts was that they all originated from type-2 *T. gondii* strains.

Intriguingly, the majority of analyzed parameters remained uniform, *i.e.* circularity, *RPC*, *FD* and *PD* did not differ among different parasite strains, time points, and hosts.

The analyzed cysts widely differed in diameter, with the cysts obtained from the experiment being smaller, albeit not significantly, compared to historical (archived) cysts. This may be due to a bias towards larger, more interesting cysts kept in a laboratory photo archive.

According to our results, circularity was a constant parameter, with all the values very close to 1, which indicates that *T. gondii* cysts in the brain tend to keep an almost perfectly round shape at any stage, a result that confirms a general assumption.

*FD*, the value that quantifies irregularity of complex objects, can range from 1 to 2 for two-dimensional images [27]. In this study, the mean *FD* value of all 31 analyzed cysts was 1.734 with very low variation among the cysts (SD=± 0.063). Despite all the differences among the cysts, the *FD* remained stable. On the other hand, lacunarity, as a measure of heterogeneity [26], had generally moderate values (0.309 ± 0.061) but showed variation among the different parasite strains. Cysts of the BGD1 strain had significantly lower lacunarity in comparison to the three other strains, which indicates these cysts were the most homogeneous. This strain of human origin was isolated at the NRLT in 2005 and maintained in mice to date, and four out of the six analyzed images were obtained from the last, 32^nd^ passage. Whether the lacunarity value is an intrinsic feature of a particular strain or is related to the *in vivo* maintenance remains to be determined.

In the experiment performed for this study, we used the Me49 strain as a widely used reference type-2 strain. In addition to SW mice commonly used in our laboratory, we included an inbred (BALB/c) mouse strain to overcome variability associated with infection in outbred mice. Moreover, potential variability caused by poor control of the size of the infecting inoculum if infection is carried out with a specific number of cysts (containing an unknown number of bradyzoites) was overcome by using (readily countable) tachyzoites. The resulting cysts were harvested at two different time points to monitor potential differences among brain cysts over time. The results showed that, irrespective of the host genetic background and cyst age, main fractal geometry values among Me49 cysts were stable*, i.e.* no significant differences were found either for *FD* or lacunarity.

Watts et al. [7] used a different imaging software (BradyCount 1.0) to estimate the number of bradyzoites from purified Me49 cysts based on the number of nuclei in the cysts, which were precisely visualized by immunostaining. The resulting value was named “packing density”, a term corresponding to what we defined in our methodology as *RPC*. Calculation of packing density included the watershed algorithm used for particle analysis in our study. The packing density in the Watts study tended to decrease with the cyst diameter; such a trend was also observed in our study for *RPC*. The simplified methodology used here, not requiring additional manipulation of brain tissue, provided comparable results with the above mentioned study, indicating the potential of computational image analysis.

This study is the first application of fractal analysis in describing the structural complexity of *T. gondii* cysts. The presented results further our insight into some computable parameters of cyst morphology. Since this study was limited to type-2 strains, it would be interesting to examine cysts of other *T. gondii* genotypes. The methodology used here is straightforward and cost-effective, requiring basic steps that are routinely applied in *T. gondii* bioassays. A practical application may include discriminating *T. gondii* cysts from similar structures including artefacts, thereby improving the specificity of bioassays [28, 29]. Moreover, as particle analysis allows for precise cell counting, it may be applied as a low cost alternative to immunostaining and other elaborate high-cost techniques used for these purposes.

## Supporting information

**S1 Data - Morphological and particle analysis parameters of tissue cysts (n=31).**

**S2 Data - Fractal dimension and lacunarity of tissue cysts (n=31).**

**S3 Data - Binary images of *T. gondii* cysts segmented by watershed algorithm (n=31).**

